# Mating duration and prolonged post-copulatory associations in *Arrenurus* water mites (Actinotrichida: Parasitengonina: Hydrachnidiae)

**DOI:** 10.1101/2021.05.23.445345

**Authors:** Mariusz Więcek, Jacek Dabert, Heather Proctor

## Abstract

Conflicts of interest between the sexes over control of mating can be reflected in various aspects of morphology and behaviour, including structure of genitalia and copulation duration. In *Arrenurus* water mites (Hydrachnidia: Arrenuridae) there are two main patterns of sperm transfer that differ in degree of potential male control of sperm uptake by females. In some species, males are equipped with an intromittent organ (the ‘petiole’) that is used to forcefully insert sperm into the female reproductive tract. In others, males lack an intromittent organ and females appear to push sperm into their reproductive opening themselves. Theory suggests that the amount of time spent in courtship after sperm transfer should differ between males with and without an intromittent structure. We predicted that male *Arrenurus* able to push sperm into the female’s reproductive tract (petiolate males) should spend less time courting females after transferring sperm than apetiolate males, which may have to ‘convince’ females to take up their sperm. Here, we examined durations of mating for 10 species of *Arrenurus* with males that differ in genital morphology: six species with males equipped with a well developed petiole (= ‘petiolate’ species) and four species with males that either completely lack a petiole or have a minute peg-like petiole that does not appear to function as an intromittent organ (= ‘apetiolate’ species). We tested whether males of petiolate species spend less time in the stage of courtship that takes place after sperm transfer (= ‘post-transfer courtship’) than apetiolate males. In contrast to our prediction, we found that species with well developed petioles spent significantly more time in post-transfer behaviours than species lacking petioles. The possible function of protracted post-transfer courtship in the genus *Arrenurus* is discussed.

## 1. Introduction

Sexual selection theory was initially proposed to explain the existence of sexual dimorphism in species in which there is little ecological differentiation between the sexes (Darwin 1859). The originally described scenarios relate to situations in which males compete for access to females through combat or through courtship displays in which females choose the winner (Darwin 1871). However, there are situations where males show morphological, physiological and behavioural adaptations that appear to bypass female choice. Sexual conflict theory argues that in some species, male genitalia and pre-copulatory behaviour evolved to force females to mate (= coercion); if this reduces female fitness, there is then selection for female resistance to male coercive morphology and behaviour (Arnqvist & Rowe 2005). Evolutionary conflicts of interest can apply to which sex controls mating rate, copulation duration, female remating propensity and ultimately fertilization of eggs (Edvardsson & Canal 2006).

Similarly, although females may benefit from mating with additional ‘higher quality’ males after a first mating, it is in the first male’s interest to prevent females from remating (Arnqvist & Rowe 2005; Firman et al. 2017). There are various adaptations to reduce polyandry which may take the form of using mating plugs by males, transferring seminal toxins or postcopulatory mate-guarding (Arnqvist & Rowe 2005; Firman et al. 2017). It has been experimentally shown for some species that prolonging the postcopulatory stage of mating increases probability of fertilization of eggs by sperm of a particular male, since more sperm and accessory ejaculate substances is transferred over time (Eberhardt 1985; Arnqvist & Rowe 2005). In response to male adaptations to prolong mating duration, females evolved counter-adaptations such as struggling in order to dislodge males (Edvardsson & Canal 2006; Bergsten & Miller 2007).

Sperm-transfer behaviour in water mites of the genus *Arrenurus* Dugès, 1834 (Actinotrichida: Parasitengonina: Hydrachnidiae) ranges from low to high complexity among around 1000 described species (Smit 2020; Więcek et al. 2021). In males, the hindbody (the cauda) may be simple and only slightly differ from female’s hindbody, whereas in other species the males have elongated or complexly sculptured cauda (Fig. 1A, B). The most complex cauda are equipped with pygal lobes, protrusions and an elaborate median intromittent organ (the petiole) (Fig. 1B). The male’s cauda and more anterior region of the hindbody has pairs of glandularia that produce an adhesive secretion during mating that is used to glue the female to the male’s hindbody (Jin et al. 1997). Moreover, males of some species have a spur-like extension on the fourth leg that is used to clasp the female’s legs and held her during the early stages of pairing (Jin et al. 1997; Proctor & Wilkinson 2001).

**Figure 1.**
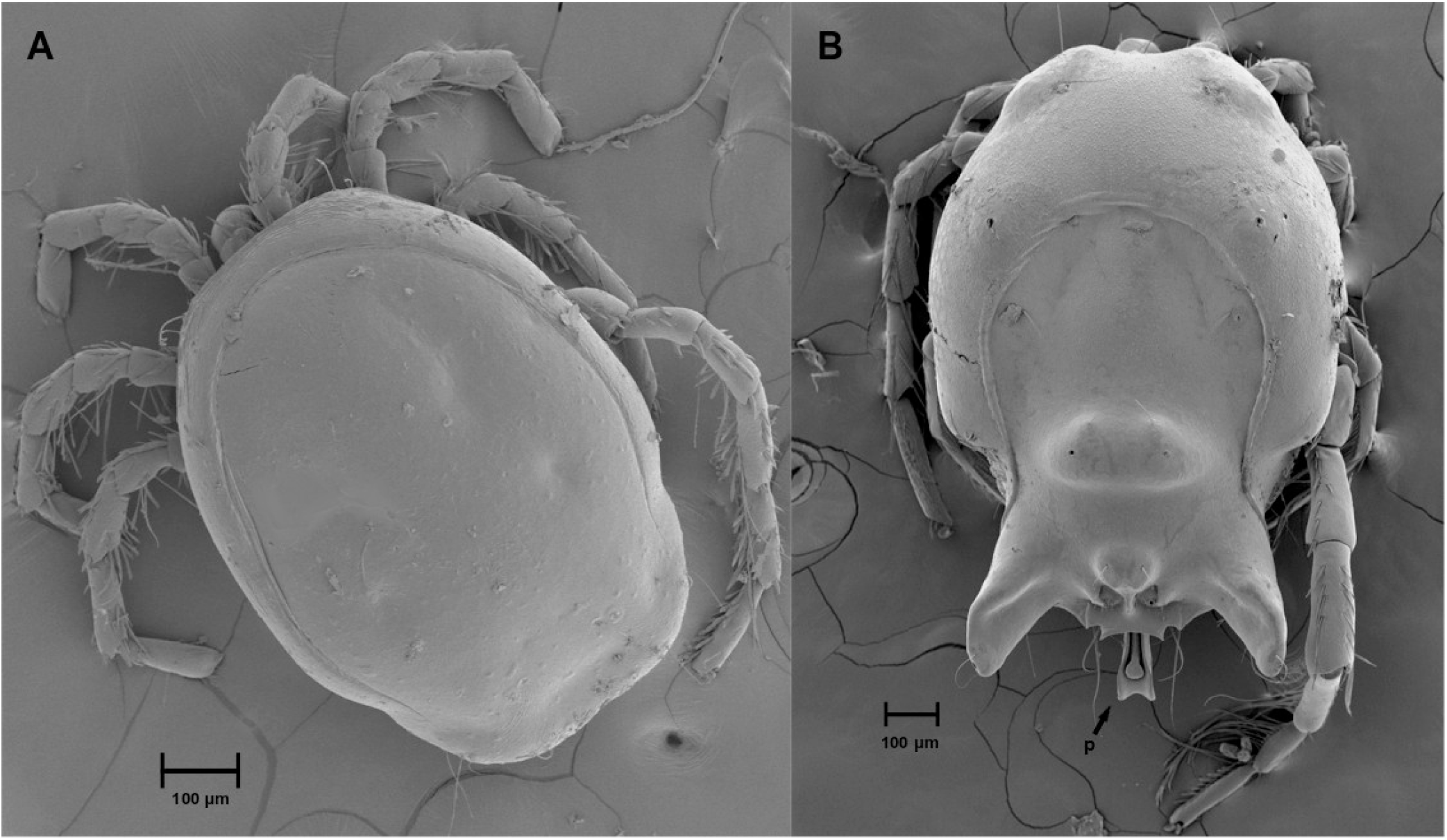
Scanning electron micrographs of males of apetiolate *Arrenurus fontinalis* (B) and petiolate *Arrenurus bicuspidator* (B); the petiole – p, indicated with an arrow.

Among *Arrenurus* there are two main modes of sperm transfer that appear to differ in the amount of control the female has over moving sperm into her genital tract. In one mode, males forcefully push sperm into the female’s genital tract with the use of well developed intromittent organ (e.g., *Arrenurus valdiviensis* K.O. Viets, Böttger 1965). In the second mode, males lack an intromittent organ. Males of these species deposit spermatophores on the substrate and lower the female on top of the sperm packets on the spermatophores. Females may have their genital flaps open at this point, or they may be closed, and they later use their legs to move the accumulated sperm from their external genital flaps into their genital openings, as in *Arrenurus globator* (O.F. Müller) (Lundblad 1929; Böttger 1965) and *A. manubriator* Marshall (Proctor 2003). Sexual conflict is likely more intense in species with males possessing well developed intromittent organs that appear to bypass female choice of sperm uptake, while female choice should be the stronger force of selection in species with males not equipped with such an organ (Proctor & Smith 1994; Proctor & Wilkinson 2001).

Both male and female *Arrenurus* are known to mate repeatedly in their life span and therefore potential conflicts over mating rates and courtship duration may take place (Proctor & Wilkinson 2001; Proctor 2003). The duration of mating in different *Arrenurus* species seems to be largely controlled by males through various methods: the male gluing the female to his hindbody, clasping and maneuvering the female before and during coupling with the use of a spur-like extension on the fourth leg, and (in some species) using an intromittent organ to insert sperm into the female. Separation of male and female at the end of the mating sequence also sometimes seems to be controlled by the male pushing the female in a way that breaks the binding of the glue (Proctor & Wilkinson 2001). Here, we observed full mating sequences and examined mating durations of *Arrenurus* species differing in presence of an intromittent structure (the petiole) in the male. We predicted that apetiolate males should spend more time on post-deposition courtship in order to ‘encourage’ females to move sperm higher into their genital tracts. In contrast, we predicted that petiolate males equipped with a well developed intromittent organ that forcefully pushes sperm into the female genital tract should spend less time on courtship following sperm transfer.

## 2. Materials and methods

### 2.1. Mite collection and maintenance

*Arrenurus* mites were collected using a net with a mesh size of 250 µm in four locations in Poland: ponds at Adam Mickiewicz University campus in Poznań (52°27’59.3”N 16°55’57.2”E), Bagnisko lake (53°29’55.7”N 16°28’42.3”E), Łagowo lake (52°19’33.6”N 15°17’16.8”E) and an unnamed pond near Zielona Góra (51°55’03.8”N 15°28’50.6”E). Data for species originating from Canada were obtained from Proctor and Wilkinson (2001). Live specimens were sorted under a stereoscope microscope in the laboratory and determined to species using a key in Gerecke et al. (2016).

Deutonymphs collected in the field were maintened in its own wells with 2 cm in diameter and 1 cm deep until they transformed into an adult. Adult mites sampled from ponds were separated by species and sex, and maintained together in the laboratory in tissue culture plates with diameter 5 cm and 3 cm deep at ambient temperatures (∼20-25 °C). Both adults and deutonymphs were fed with ostracods, cladocerans and copepods from laboratory colonies established based on zoological material collected in local water bodies.

### 2.2. Mating observations

Female and male mites (whenever possible, virgin specimens) were held individually for 24 h before observation in microaquaria that were half filled with tap water. The well-plates had diameter of 2 cm and depth of 1 cm with the scratched bottom to create a stable substrate for the mites during mating. We took videos of the entire mating sequences of seven species of *Arrenurus*: five with petiolate males (*A. bicuspidator, A. bruzelii, A. claviger, A. cuspidator, A. tricuspidator*) and two with apetiolate males (*A. globator* and *A. stecki*). Details of mating behaviour of these species are in Więcek (2016). Video recording began when a male was introduced to a container with a conspecific female. Recording was terminated upon physical separation of individuals after mating. In order to obtain a larger sample size to test our predictions, the results were pooled with the data obtained by Proctor and Wilkinson (2001) for three North American species: petiolate *A*. nr. *reflexus* and apetiolate *A. manubriator* and *A. rufopyriformis*. The observations of mating behaviour were made with Zeiss Stemi 2000-C stereomicroscope and videotyped with the use of attached to the stereomicroscope DIC illumination and digital camera Olympus DP71 with CellD 2.8 software (Olympus Soft Imaging Solutions GmbH).

We divided mating sequences into the following categories, listed in temporal order: pre-pairing stage, behaviours from initial pairing to the end of spermatophores deposition and post-deposition behaviour. In all categories except pre-pairing, the female was attached to the male. For statistical analysis we calculated the following: pre-pairing duration = total time from introduction of a male into a female’s container to the female being glued onto the male; total duration of mating = total time that male and female were attached (i.e. glued together), including time spent in pre-deposition behaviour, spermatophore deposition and sperm transfer (deposition of spermatophores was assumed based on male ‘dipping’ movements), and post-deposition behaviour; post-deposition behaviour = time male and female were still attached to each other after sperm transfer. Unlike in Lundblad (1929) and Więcek (2016), we treated the male behaviours of lateral waving and alternate bending of left and right legs as part of post-deposition behaviour.

SEM photographs of couples in copula were made in the Department of Earth and Atmospheric Sciences, University of Alberta. After dehydration through an alcohol-HMDS (hexamethyldisilazane) series, mites were mounted on stubs using double-sided tape, sputter coated with gold, and examined using a JEOL 630 I field emission scanning electron microscope (SEM) in the Department of Earth and Atmospheric Sciences, University of Alberta. SEM photographs were edited in Photoshop 6.0.

### 2.3. Statistical analyses

Data were tested for significant deviations from normal distribution with Shapiro-Wilk test. No significant deviations from normal distribution for comparing all groups in analyses were found, and therefore parametric two sample *t*-tests (two tailed) were applied. The sample sizes for statistical analyses were based on numbers of species of petiolate and apetiolate *Arrenurus* (n=6 and 4, respectively). The data on difference in percentage of time spent on post-spermatophore-transfer behaviours were changed to proportions and arcsine-transformed (Sokal & Rohlf 1995). Moreover, mean values and confidence intervals (95%) for the estimate of the mean were calculated based on the standard errors.

The duration of pre-pairing stage of mating can be affected by female age and whether they were previously exposed to males (H. Proctor, pers. obs.). Because mating status and age of mites were not standardized, pre-pairing duration was excluded from statistical tests. All analyses were run with the statistical software PAST 4.03 (Hammer et al. 2001).

## 3. Results

In this study 31 pairings of 7 *Arrenurus* species were observed (*A. bicuspidator, A. bruzelii, A. claviger, A. cuspidator, A. tricuspidator, A. globator, A. stecki*), which resulted in about 160 hours of mating trials (Table 1). Moreover, observations of *A. manubriator, A*. nr. *reflexus* and *A. rufopyriformis* were included in the statistical analyses (Proctor & Wilkinson 2001). Total duration of mating duration ranged from 29.00 ± 5.00 minutes in apetiolate *A. stecki* to 630.00 ± 79.00 minutes in petiolate *A. bicuspidator* (Table 1, SEM photographs of mites in copula are in Fig. 2). Total duration of mating in petiolate species was significantly longer (mean = 455.7 ± 55.16 min, n=6) than in apetiolate species (mean = 85.45 ± 25.32 min, n=4) (*t*-test: 6.10, two-tailed *P<*0.001) (Fig. 3A). There was no significant difference in duration of behaviours from initial pairing to the end of spermatophores deposition between petiolate and apetiolate species (*t*-test: 0.81, two-tailed *P*=0.44). Petiolate species spent a significantly higher percentage of total mating time on post-deposition behaviour (mean = 93.69± 1.06%) than apetiolate species (mean = 55.27 ± 12.47%) (*t*-test: 3.61, two-tailed *P* = 0.033) (Fig. 3B).

**Table 1.**
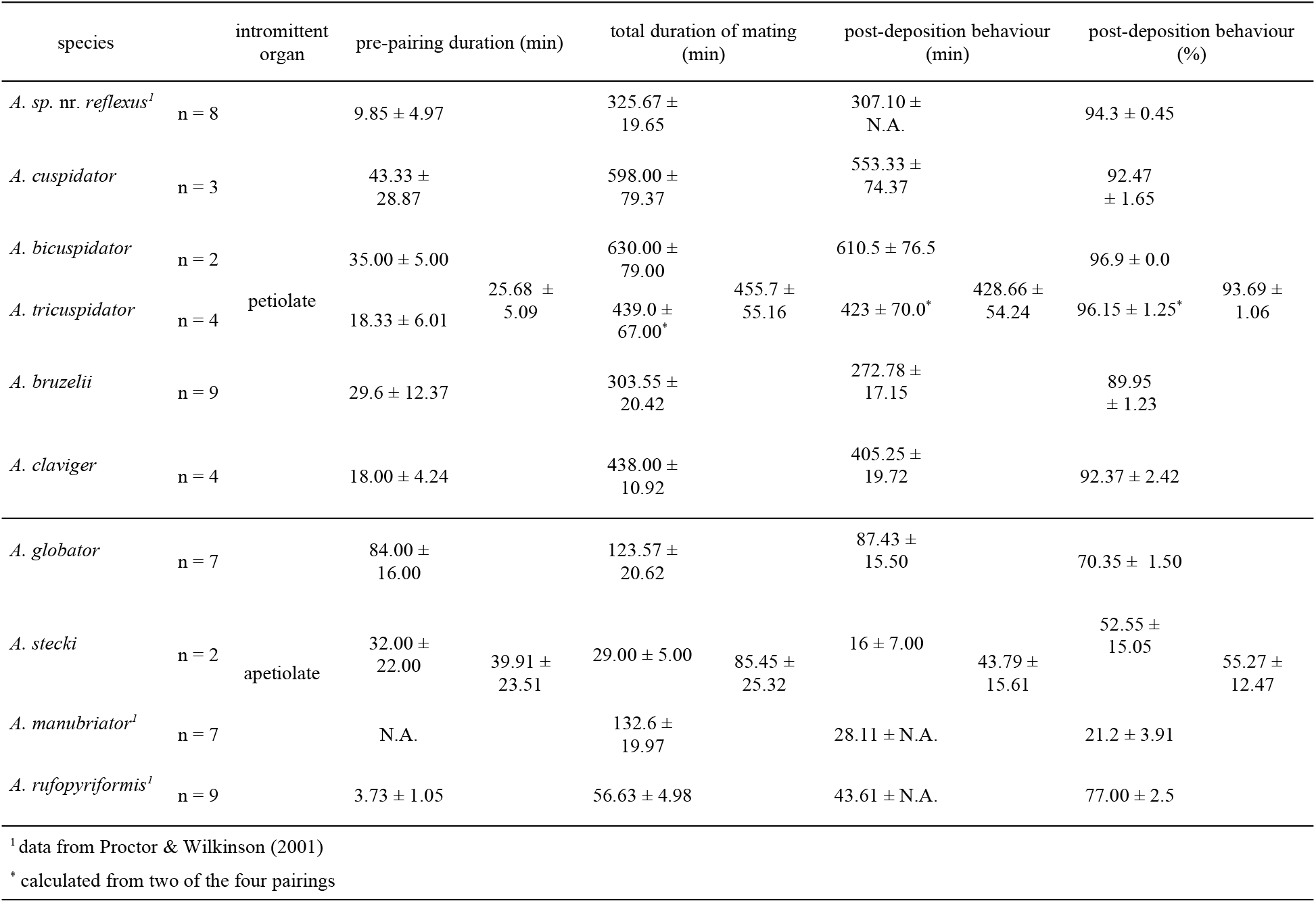
Mean time (± SE) spent on particular stages of mating in different species of the genus *Arrenurus*. The post-deposition behaviour is part of pairing behaviour and is expressed as percentage of the total time spent on mating (also given in min). The total duration of mating does not include pre-pairing stage.

**Figure 2.**
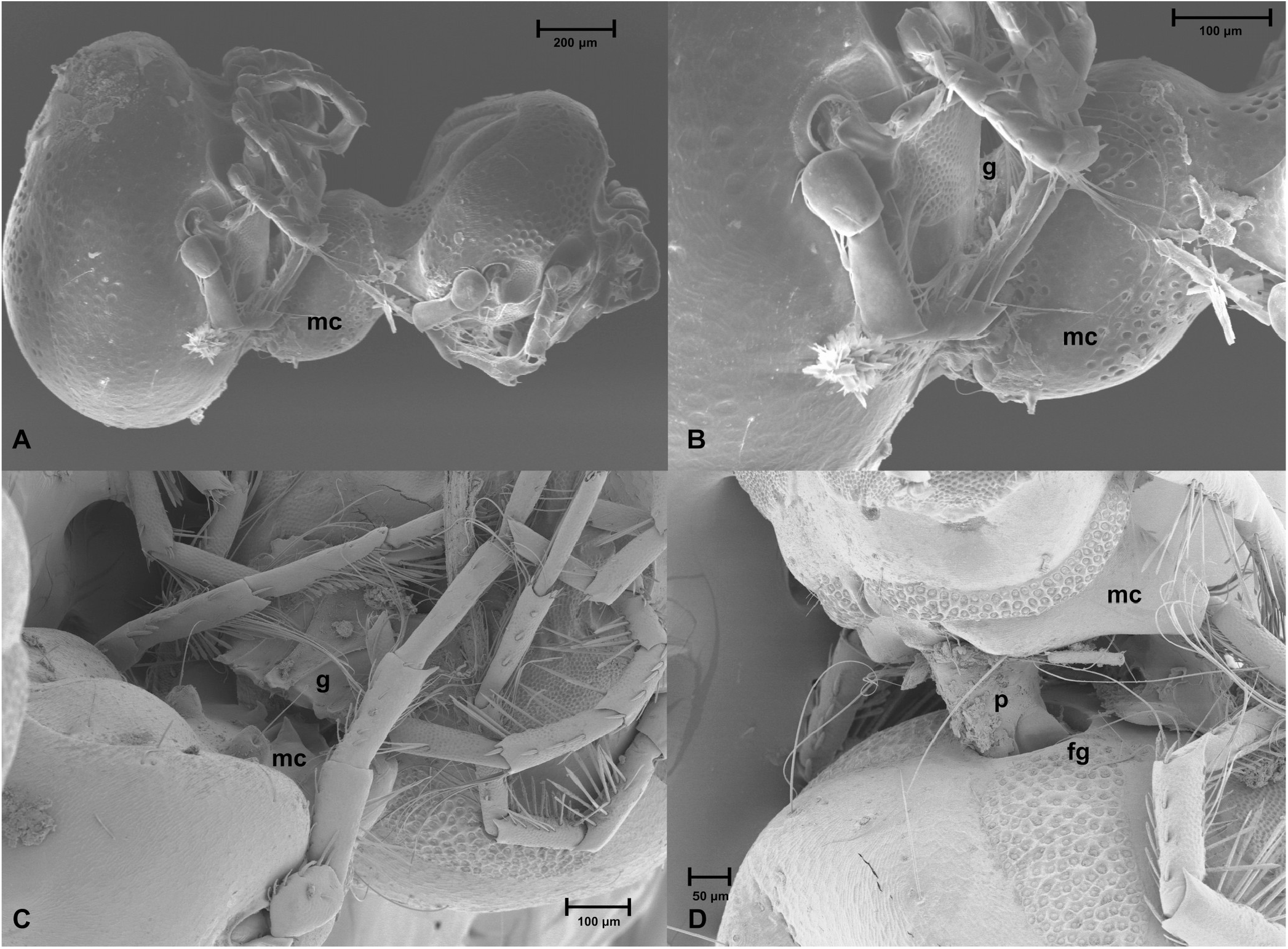
Scanning electron micrographs of *Arrenurus* spp. in copula. A, B. *Arrenurus globator* in mating position (lateral view): female glued to males hindbody (no petiole). C, D. *Arrenurus* sp.: female glued to males dorsum; the petiole inserted in to female genital opening is shown. Abbreviations: p – petiole (intromittent organ), mc – male’s cauda (hindbody), g - adhesive secretion enabling gluing female to male’s hindbody produced by glandularia, fg – female’s genital opening.

**Figure 3.**
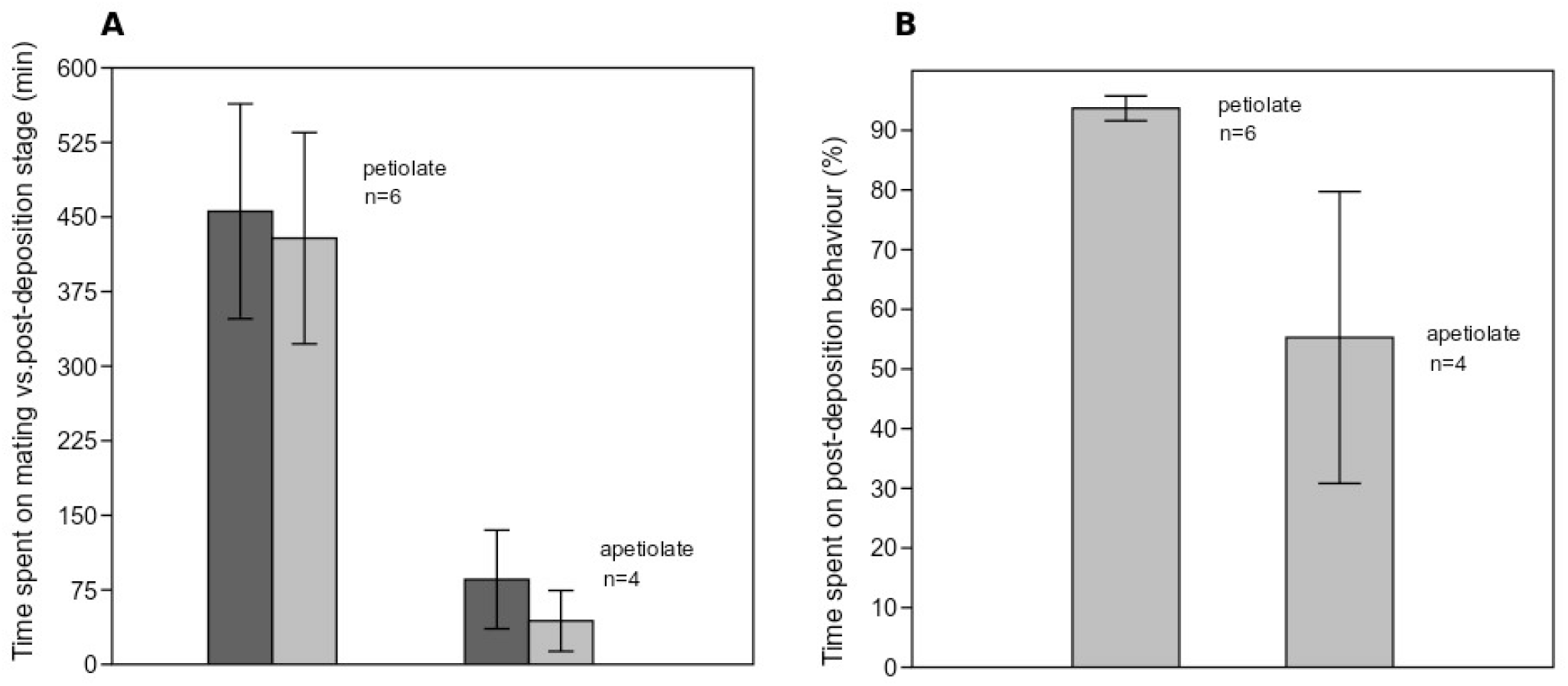
Mean time (and standard errors) spent on mating and post-deposition behaviours: **A**. Mean time spent on mating (dark grey) verus post-deposition stage (light grey) in petiolate and apetiolate species (in min). **B**. Mean time spent on post-deposition behaviours (expressed as percentage of the total time spent on mating). Sample sizes and SE are shown.

## 4. Discussion

We expected that *Arrenurus* males with an intromittent structure (the petiole) would spend less time in courtship behaviours after sperm transfer than apetiolate males, which have less coercive control over whether sperm enters the female’s genital opening. Contrary to our prediction, males of petiolate species spent more time in post-deposition behaviours than males of apetiolate species, both absolutely and relatively (as proportion of total time spent mating). Although both petiolate and apetiolate species exhibited prolonged post-deposition associations, in petiolate species post-deposition behaviour made up > 90% of total mating time in contrast to ∼50% in apetiolate species. However, there was a considerable variation in time spent on post-transfer behaviours among apetiolate species.

Why might males of petiolate species show such long periods of attachment to females after introducing sperm into her genital tract? Protracted courtship or simply male attendance following sperm transfer is known to occur in different animal groups and is thought to reduce the probability of the female to be inseminated by a subsequent male (mate guarding, Arnqvist & Rowe 2005). Also, in this stage of mating males can manipulate female’s physiology by stimulating oviposition or decreasing production of sex pheromones attracting males (Arnqvist & Rowe 2005; Mazzi et al. 2009). Radwan and Siva-Jothy (1996) demonstrated that males of the terrestrial mite *Rhizoglyphus robini* (Acaridae) increase fertilisation of eggs by their sperm by prolonging attachment to a female. We hypothesize that this could be an explanation of the function of periods of motionlessness and courtship behaviour after sperm transfer in the examined *Arrenurus* spp.

It is well known that in various invertebrate taxa the risk of sperm competition can be selective force in males and can be expressed by using mating plugs and mate guarding that are supposed to prevent future inseminations (Dougherty et al. 2016; Firman et al. 2017). Male courtship and postmating behaviours may also encourage preferential use of sperm of a particular male and thus can be under selection by cryptic female choice (Firman et al. 2017). It was shown that *Arrenurus* spp. can mate repeatedly (e.g. *A. manubriator*; Proctor 2003), which potentially leaves space for both selective pressures associated with sperm competition between males and physiological processes in females affecting preferential use of sperm of a particular male. Males of apetiolate and petiolate species may differ in the amount of sperm competition they face, if, in addition to pushing sperm into the female’s genital opening, the petiole also acts to remove or displace sperm deposited by previous males as does, e.g., the genitalia of male damselflies (Waage 1979). If so, the greater duration of both pre- and post-transfer behaviour in petiolate species may reflect the lengths to which such males must go to increase the likelihood that females will use their sperm and not that of a previous or subsequent mate. To test this idea, the paternity of egg clutches could be assessed in matings in which the second male is allowed to complete the entire mating sequence or is interrupted at various times after sperm transfer.

Finally, although optimal copulation duration can be a subject of conflict between the sexes, which may be the case in the examined *Arrenurus*, it has been observed that females can benefit from prolonged copulations, for instance, because they receive larger ejaculates (e.g. bruchid beetles, Edvardsson & Canal 2006). Therefore, further research should be done to determine if prolonged post-copulatory associations in *Arrenurus* mites influence female’s lifetime fecundity as well as male’s paternity.

## Acknowledgements

We thank Dr. G. Greczka, Poznań, Poland who provided statistical advice. The study is part of the International PhD Programme ‘From genome to phenotype: A multidisciplinary approach to functional genomics’ (MPD/2010/3) funded by the Foundation for Polish Science (FNP). Additional funding was provided by a Natural Sciences and Engineering Research Council of Canada (NSERC) Discovery Grant to HCP.

